# Mushroom bodies are required for accurate visual navigation in ants

**DOI:** 10.1101/2020.05.13.094300

**Authors:** Cornelia Buehlmann, Beata Wozniak, Roman Goulard, Barbara Webb, Paul Graham, Jeremy Niven

**Affiliations:** School of Life Sciences, University of Sussex, Brighton BN1 9QG, UK; School of Informatics, University of Edinburgh, Edinburgh EH8 9AB, UK

**Keywords:** Visual navigation, innate visual behaviour, neural circuitry, mushroom body, brain lesions, procaine hydrochloride, wood ants, *Formica rufa*, insects

## Abstract

Visual navigation in ants has long been a focus of experimental study [1–3], but only recently have explicit hypotheses about the underlying neural circuitry been proposed [4]. Indirect evidence suggests the mushroom bodies (MB), a known site of olfactory learning [5–10], may also be the substrate for visual memory in navigation tasks [11–14]. Computational modelling shows that MB neural architecture could support this function [15, 16], though there is no direct evidence that ants require MBs for visual navigation. Here we show that lesions of MB calyces impair ants’ visual navigation to a remembered food location whilst leaving their innate responses to visual cues unaffected. Ants are innately attracted to a large visual cue but we trained them to locate a food source at a specific angle to this visual cue. Subsequent bilateral or unilateral lesioning (through procaine hydrochloride injection) of the MB calyces, caused ants to revert to their innate cue attraction whilst control (saline) injected ants still approached the feeder. The ants’ path straightness and walking speed were unaffected by lesions. Reversion towards the cue direction occurred irrespective of whether it was ipsi-or contralateral to the lesion site, showing this is not due simply to an induced motor bias. Monocular occlusion did not diminish ants’ ability to locate the feeder, suggesting the lesion is not merely interrupting visual input to the calyx. The demonstrated dissociation between innate and learnt visual responses provides direct evidence for a specific role of the MB in navigational memory.

## Results And Discussion

### Ants learn to navigate to a food source relative to a visual cue

Wood ant (*Formica rufa*) foragers were placed at the centre of a circular platform within a large, circular white arena (Figure 1A). A large black rectangular cue was mounted on the wall of the arena. Foragers were innately attracted to the cue, walking towards it (Figure S1A). We then trained foragers to find food located at the edge of a circular platform placed 30° to the right of the cue (Figure 1B). After 3.8 ± 1.1 days (mean ± SD) days of training, the ants’ direction relative to the cue had shifted significantly (Watson Williams test, p<0.001) and they walked directly towards the feeder (Figure S1B). This demonstrates that ants can visually navigate to a food source, overriding their innate attraction to the visual cue.

**Figure 1:**
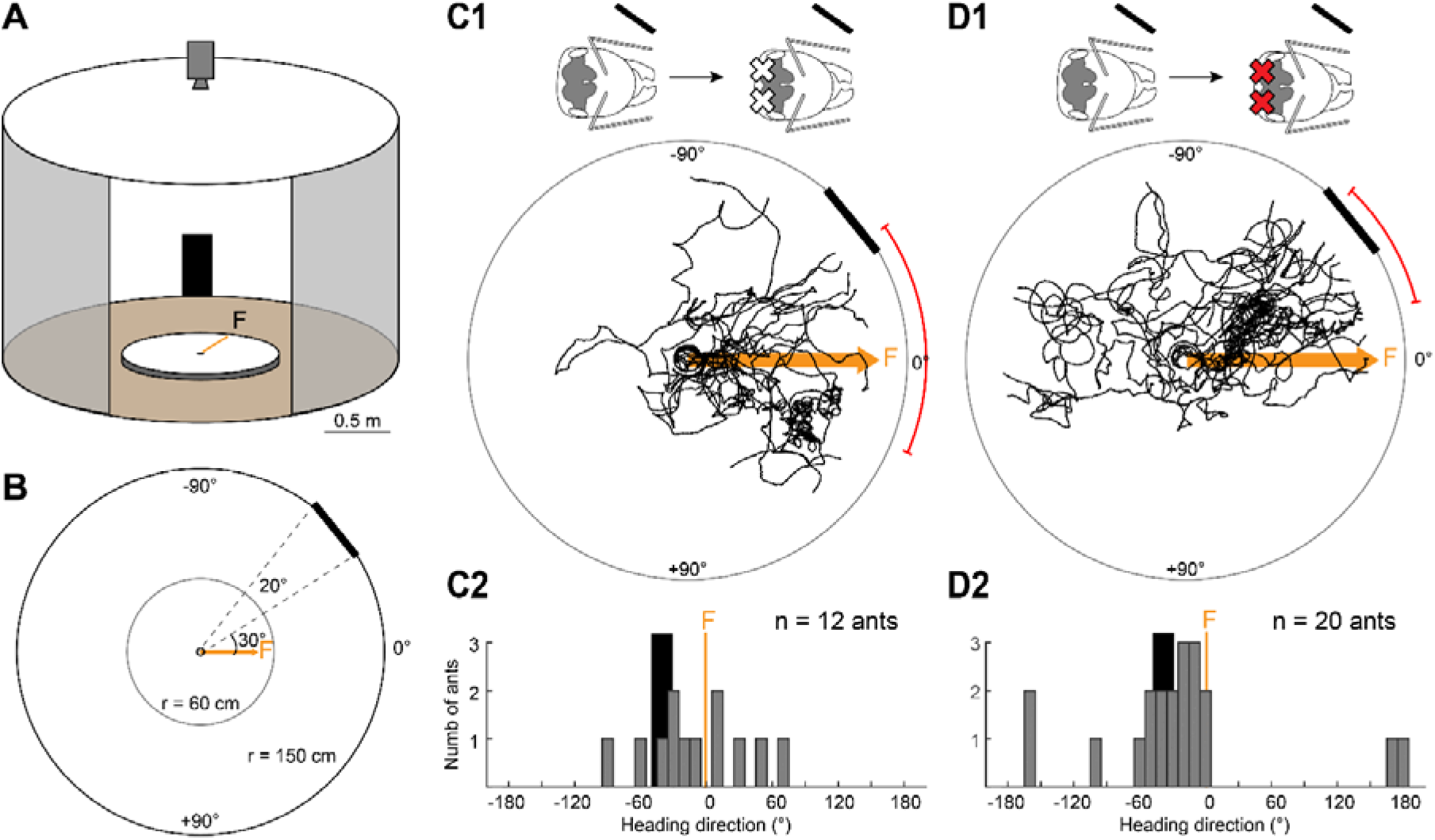
Mushroom bodies (MBs) are required for accurate visual navigation. **(A)** Experimental setup: Wood ants learnt to navigate to a feeder (F) at the edge of a circular white platform (radius: 60 cm). F was placed 30° to the right edge of a 20° wide black rectangle (height: 90 cm, width: 52 cm) mounted at the inner wall of the surrounding cylinder (radius: 1.5 m, height: 1.8 m). A camera recorded the ants’ path from above. A small door allowed us to access the arena but for clarity the door is shown here open and larger. For details see STAR Methods. **(B)** Behavioural setup shown in (A) depicted from the top. **(C)** Ants were trained to the feeder (F) in the behavioural arena shown in (A). Once ants were well-trained, we performed a bilateral chemical injection in the MB calyces and recorded the navigational behaviour of the injected test ants. **C1**: Control solution (rhodamine in saline) was injected bilaterally into the MB calyces. Injection sites are shown as X in the schematic ant-heads, and the black bar to the side of the ant’s head represents the side on which the visual cue would appear as the ant progressed to the feeder. Paths of individual control ants are shown as black paths. Bootstrapped 95% confidence interval (CI) for the medians of the heading directions is shown as red arc. Orange arrow shows the direct way to the feeder (F). For simplicity the visual cue is shown here at the edge of the platform (r = 60 cm) instead of its true position at the cylinder wall. **C2**: Heading directions of control ants at r = 50 cm. Black rectangle represents visual cue. Feeder (F) is shown in orange. Bin size, 10°. **(D)** Same as (C) but for lesioned ants with the local anaesthetic (procaine hydrochloride and rhodamine in saline) injected bilaterally into the MB calyces.

### Mushroom bodies (MBs) are required for accurate visual navigation

We investigated the role of the ants’ mushroom bodies during this visual navigation task by making chemical lesions through the injection of procaine hydrochloride, a local anaesthetic that silences neural activity by reversibly blocking voltage-gated channels, including voltage-gated Na^+^ channels [11, 17–19]. The MBs are bilaterally paired structures in the insect brain that receive direct visual input from the optical lobes to the visual input region in the calyces [20–26]. Hence, we lesioned both calyces of individual ants, co-injecting a fluorescent dye (rhodamine) to ensure these lesions were correctly targeted (see STAR Methods; Figure S1E1). As a control, we injected ants with rhodamine in saline.

After the injection, ants were returned to the arena for testing. Both control and lesioned ants were still directed (Rayleigh test, p<0.001). In the control group, the learnt feeder location (0°) was within the 95% confidence interval (CI) of the ants’ heading directions (bootstrap distribution of the median; CI: [-34.4° (2.5 percentile values), 22.9° (97.5 percentile value)]). Hence, control ants were still directed towards the feeder (Figure 1C). For lesioned ants, however, the feeder location was outside the paths’ 95% CI ([-45.9°, −14.0°]), showing that lesioned ants no longer approached the learnt feeder location (Figure 1D). Thus, bilateral silencing of the MB calyces through chemical lesions caused ants that had learnt to visually navigate to a food source to revert towards their innate attraction to the visual cue. Moreover, bilateral silencing did not significantly alter the ants’ walking speed or path straightness (Kruskal Wallis test with Dunn’s post hoc and Bonferroni corrections, p>0.05; Figure S2) relative to the control ants, suggesting that the deficit is specific to the learnt visual navigation behaviour.

### Unilateral lesions in the MB calyces affects learnt visual navigation but not innate behaviour

Reversion to the innate cue attraction following a bilateral lesion of the MB calyces does not exclude the possibility that a single MB calyx is sufficient for visual navigation. To test this, we trained ants as before but made a unilateral lesion, ipsilateral to the apparent position of the visual cue (Figure S1E2). Paths from control and lesioned ants were still directed (Rayleigh test, p<0.001). For the control ants, the learnt feeder location (0°) was within the 95% confidence interval (CI) of the ants’ heading directions (CI: [−18.4°, 1.5°]). Hence, control ants still navigated to the learnt feeder location (Figure 2A). In unilaterally lesioned ants, however, the feeder was outside the paths’ 95% CI ([−25.5°, −3.5°]; Figure 2B). Following a unilateral lesion there was no significant difference in the walking speed or path straightness between control and lesioned ants (Kruskal Wallis test with Dunn’s post hoc and Bonferroni corrections, p>0.05; Figure S2). These results suggest that a unilateral MB calyx lesion is sufficient to impair visual navigation causing ants that had learnt the location of a food source to revert partially towards their innate attraction to the visual cue.

**Figure 2:**
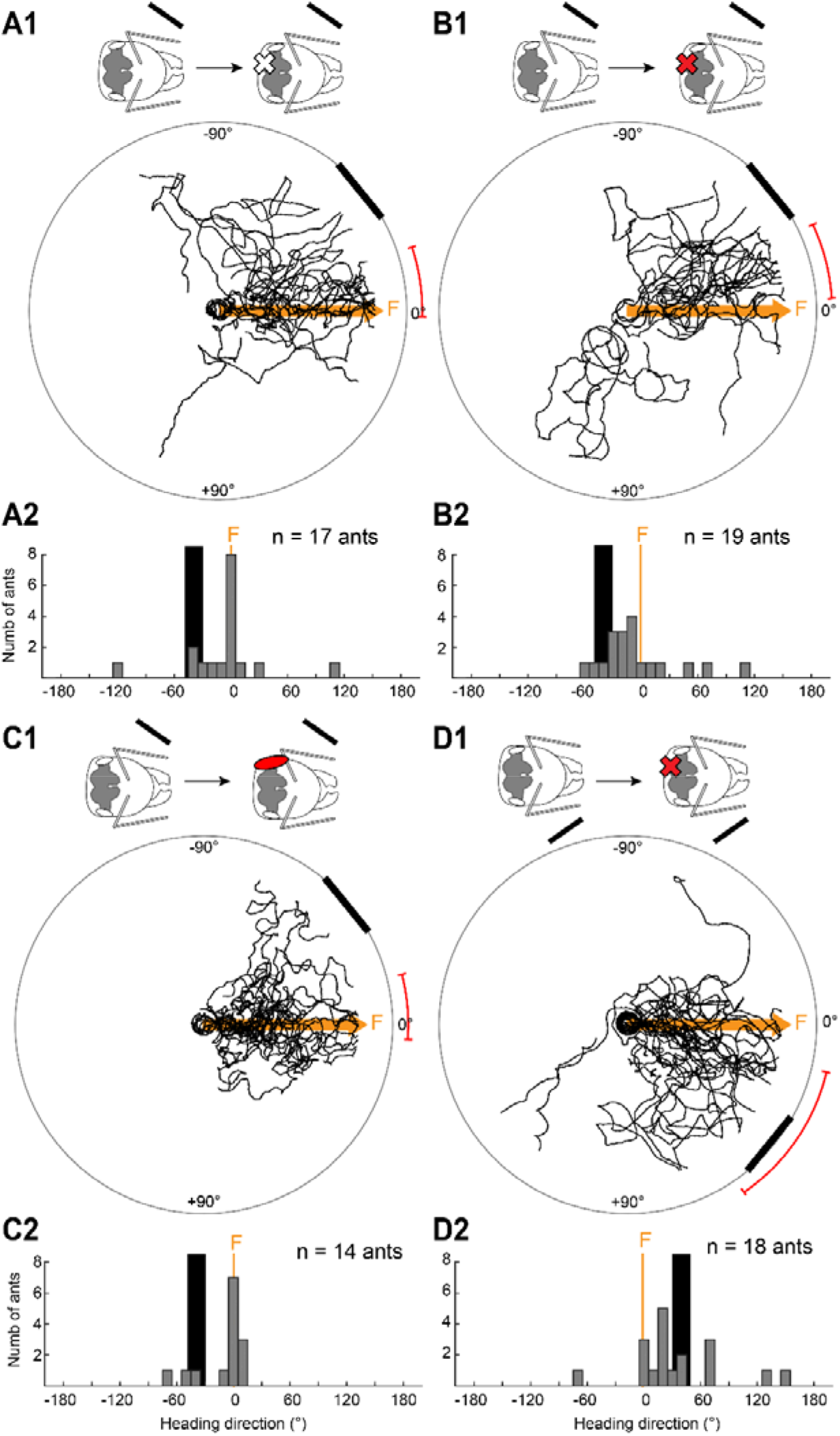
Both MB calyces are necessary for accurate visual navigation. **(A-C)** Ants were trained to the feeder (F) in the behavioural arena shown in Figure 1A. **(A)** Well-trained ants had unilaterally injections of the control solution into the MB calyx ipsilateral to the visual cue (X, injection site; black bar, apparent position of visual cue). For path plot and histogram description see Figure 1. **(B)** Same experimental procedure as in (A) but these ants had the local anaesthetic injected into the MB calyx ipsilateral to the visual cue. **(C)** Same experimental procedure as in (A) but these ants had their compound eye ipsilateral to the visual cue occluded in the test. **(D)** Ants were trained to a feeder placed 30° left of the visual cue (see Figure S1D and STAR Methods). The unilateral MB calyx lesion was contralateral to the position of the visual cue.

In contrast, the innate behaviour of lesioned naïve ants that had not previously experienced the arena or the visual cue was unaffected by unilateral MB calyx lesions (Figure S3; see also STAR Methods). The paths of innate lesioned and control ants were directed (Rayleigh test, p<0.001), and the visual cue laid within the paths’ 95% CI for both of the groups (Control ants, CI: [−43.0°, −29.0°], Figure S3A; lesioned ants, CI: [−57.7°, −36.6°], Figure S3B). Moreover, there was no significant difference in the ants’ walking speed or their path straightness (Kruskal Wallis test with Dunn’s post hoc and Bonferroni corrections, p>0.05; Figure S2). Hence, the innate attraction of wood ants to dark visual cues [27] is unaffected by lesioning a MB calyx, suggesting that there is a dissociation in brain regions required for innate and learnt visual guidance.

### The effects of MB lesions are not equivalent to an absence of peripheral visual input

The MB calyces, to which we targeted the chemical lesions, receive inputs from the visual system in ants [24–26]. To ensure that the deficit in visual navigation to the learnt feeder location caused by the unilateral lesions could not be explained by a deficit or asymmetry in visual inputs, we monocularly occluded trained ants (see STAR Methods). Paths of the monocularly occluded ants were directed (Rayleigh test, p<0.001) and the feeder was within the paths’ 95% CI (CI: [−14.4°, 4.1°], Figure 2C). Thus, the change in path direction to the visual cue produced by unilateral lesions was not caused by a deficit or asymmetry in visual input but rather, other deficits, caused by the lesion of the MB calyx.

### An induced motor bias cannot account for the shift of ants’ paths towards the visual cue

Insects are known to be lateralised in many behaviours including some aspects of learning and memory (reviewed in [28]), and wood ants are no exception showing lateralisation in behaviour [29] and memory formation [30]. To ensure that the behavioural shift towards the visual cue following unilateral lesions were not produced by a lateralised motor bias, we trained a further cohort of ants with the feeder placed 30° left of the visual cue (Figure S1D). For this cohort of ants, the unilateral MB calyx lesion was contralateral to the position of the visual cue. Lesioned ants were still directed (Rayleigh test, p<0.001) but the feeder was outside the paths’ 95% CI (CI: [14.0°, 55.7°], Figure 2D). Hence, ants no longer navigated towards the learnt feeder location. Lesioned ants’ walking speed and path straightness were not different from those of control ants (Kruskal Wallis test with Dunn’s post hoc and Bonferroni corrections, p>0.05; Figure S2). Hence, the observed shift in the ants’ heading direction after a unilateral lesion from the food source towards the visual cue cannot be accounted for by a lateralised motor bias and is instead a reversion from the learnt location to the innate visual cue attraction. However, it remains possible that the relative position (ipsi-or contralateral to the visual cue) of the lesion may affect the extent of the shift back towards the visual cue.

### Is visual guidance information transferred to the contralateral brain hemisphere?

Previous experiments have suggested that there is no interocular transfer in visually navigating ants [12, 31] strongly suggesting that visual information is used retinotopically for navigation. Furthermore, neurophysiological data has shown that ants have bilateral projections from the optic lobes to the mushroom bodies of both hemispheres [25], however, it is unclear whether, and to what extent, retinotopically organised visual information for navigation is transferred to the contralateral brain hemisphere. To address this, we monocularly occluded the compound eye contralateral to the visual cue before training commenced, thereby restricting learnt views of the visual cue to the ipsilateral eye. After training and prior to testing, we lesioned the MB calyx ipsilateral to the visual cue (Figure 3). Ants with one eye covered performed as well as normally sighted ants in the training (Watson-Williams test, p>0.05; Figure S1BC). In tests, both control and lesioned ants were still directed (Rayleigh test, p<0.001). The learnt feeder location was within the 95% CI of the heading directions of the control ants’ paths (CI: [−24.0°, 6.3°]) showing that they still navigated to this location (Figure 3A). In unilaterally lesioned ants, however, the feeder location was outside the paths’ 95% CI ([−27.2°, −9.1°]; Figure 3B) showing that in monocularly occluded ants, a unilateral lesion is sufficient to prevent visual navigation and cause reversion to innate attraction to the visual cue. Again, following a unilateral lesion there was no significant difference in the ants’ walking speed or their path straightness (Kruskal Wallis test with Dunn’s post hoc and Bonferroni corrections, p>0.05; Figure S2).

**Figure 3:**
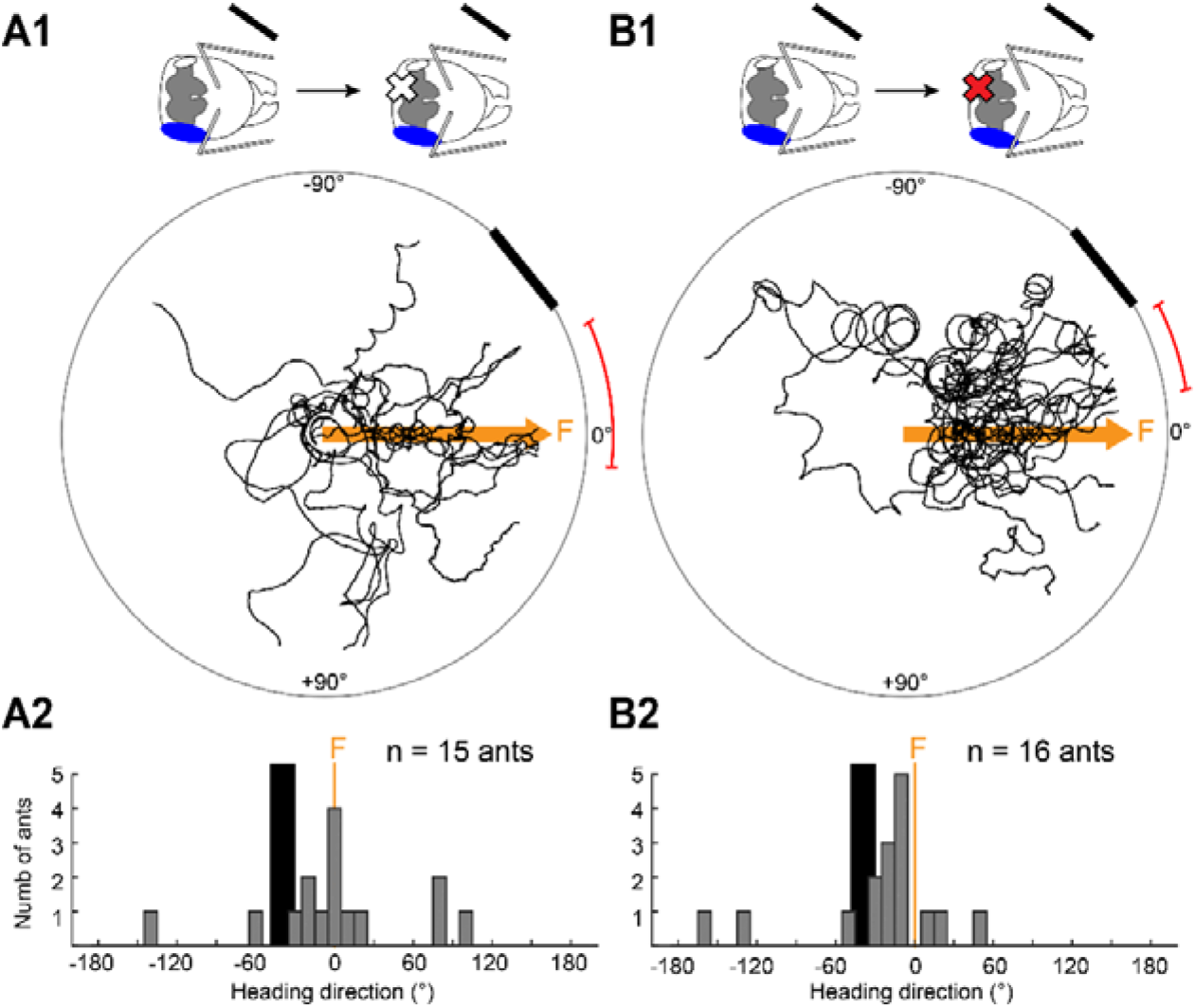
A unilateral lesion is sufficient to prevent visual navigation in monocularly occluded ants. **(A-B)** The compound eye contralateral to the visual cue was monocularly occluded before training, hence, throughout training and testing, ants received visual input only on the side ipsilateral to the visual cue. **(A)** The MB calyx ipsilateral to the visual cue was injected with the control solution, after training and prior to testing. **(B)** Same as (A) but local anaesthetic was injected on the side ipsilateral to the visual cue. For path plot and histogram descriptions see Figure 1.

### The role of the MB in visual navigation

We have provided direct evidence that the MB calyces are involved in visual navigation to learnt locations in ants. This is consistent with previous observations: the MB of ants [24–26] and other Hymenopterans [22, 23] receive direct inputs from the optic lobes; the MBs of ants [32, 33], again like those of other Hymenopterans [34, 35], expand at the onset of foraging; changes in the expression in the MB of a gene associated with learning co-occur with orientation flights in novel environments in honeybees [36]. Computational modelling has shown that the neural circuitry of the MBs is well suited for the storage of navigationally relevant visual information [4, 15, 16], specifically to encode the familiarity of multiple views. A study on cockroaches implicated the MBs in a visual place learning task but not a simple beacon aiming task [13], similar to the dissociation we observe between learnt and innate behaviour in the current study. However, in fruit flies, a place learning task [14], and some other visual learning paradigms [14, 37, 38] appear to depend on the central complex; although the MB have been shown to have a role for some visual associations [21, 39]. This suggests that the locus of visual memories within the insect brain is likely task dependent or may differ between insects from different orders. The innate visual behaviour of attraction to a conspicuous cue was not affected by MB lesions in our experiments, providing further evidence that different visual tasks involve different neural pathways.

The results we have presented are consistent with models [4, 15, 16] that suggest visually driven activity across the population of Kenyon cells in the MBs can efficiently represent experienced views, learnt through dopaminergic reinforcement of the connections between KCs and MB output neurons conveying valence [40, 41] to subsequently guide forward movements or turns. Our results suggest that each MB stores views encompassing the whole visual field, as unilateral lesions affect the behaviour in a consistent way (a reversion towards innate attraction to the cue) irrespective of the ipsi- or contra-lateral location of the cue or blocking of half the visual field during learning. The reversion appears more complete for bilateral lesions, but also for lesions contralateral to the cue, suggesting there may be overall redundancy, and perhaps also some lateralisation of function. It remains an open question how mushroom body outputs are translated into steering information for downstream motor control, although several theoretical mechanisms involving output to the central complex have been proposed [42, 43].

---

**Figure S1:**
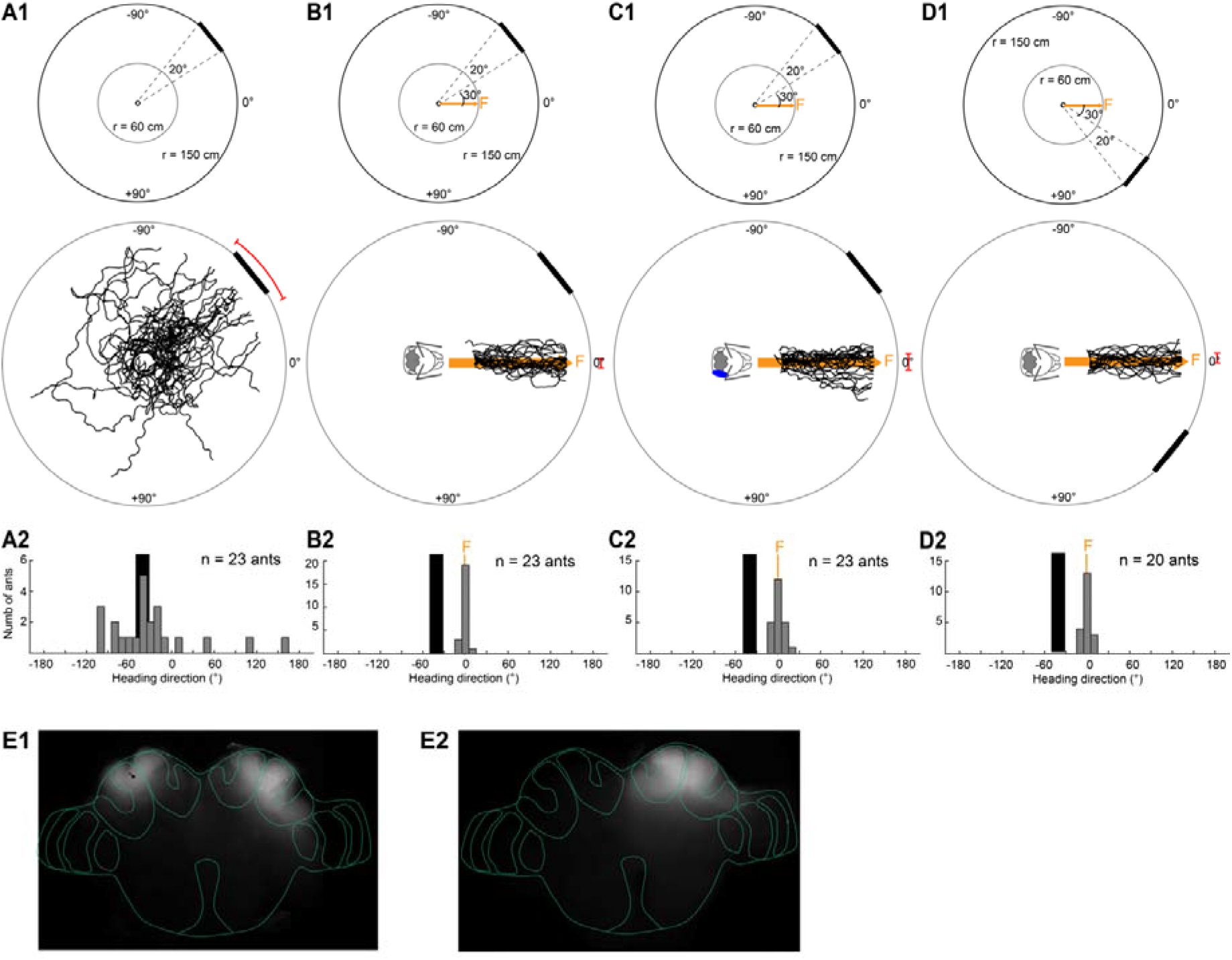
Innate and learnt visual behaviour and physiological manipulations. **(A)** Innate response of wood ants to the visual cue (shown as black bar). **A1:** Top: The ants’ paths were recorded on a circular white platform (radius: 60 cm). A 20° wide black rectangle (height: 90 cm, width: 52 cm) mounted at the inner wall of the surrounding cylinder (radius: 1.5 m, height: 1.8 m). A camera recorded the ants’ path from above. Bottom: Paths of individual naive ants are shown as black paths. For simplicity the visual cue is shown here at the edge of the platform (r = 60 cm) instead of its true position at the cylinder wall. For details see Figure 1A and STAR Methods. **A2:** Heading directions of naive ants at r = 50 cm. Black rectangle represents visual cue. Bin size, 10°. **(B)** Well-trained ants that have learnt to navigate to a feeder (F) at the edge of the circular platform. F was placed 30° to the right edge of the visual cue. For details see Figure 1A and STAR Methods. Orange arrow shows the direct way to the feeder (F). Once ants were well-trained, we performed a bilateral or unilateral chemical injection in the MB calyces and recorded the navigational behaviour of the injected test ants. **(C)** Same procedures as in (B) but these ants learnt to navigate to the feeder (F) with a compound eye unilaterally occluded throughout the training. **(D)** Same procedures as in (B) but this cohort of ants was trained to a feeder placed 30° to the left edge of the visual cue. **(E)** Physiological manipulations. **E1:** Bilateral chemical injection into the MB calyces. **E2:** Unilateral chemical injection into the MB calyx.

**Figure S2:**
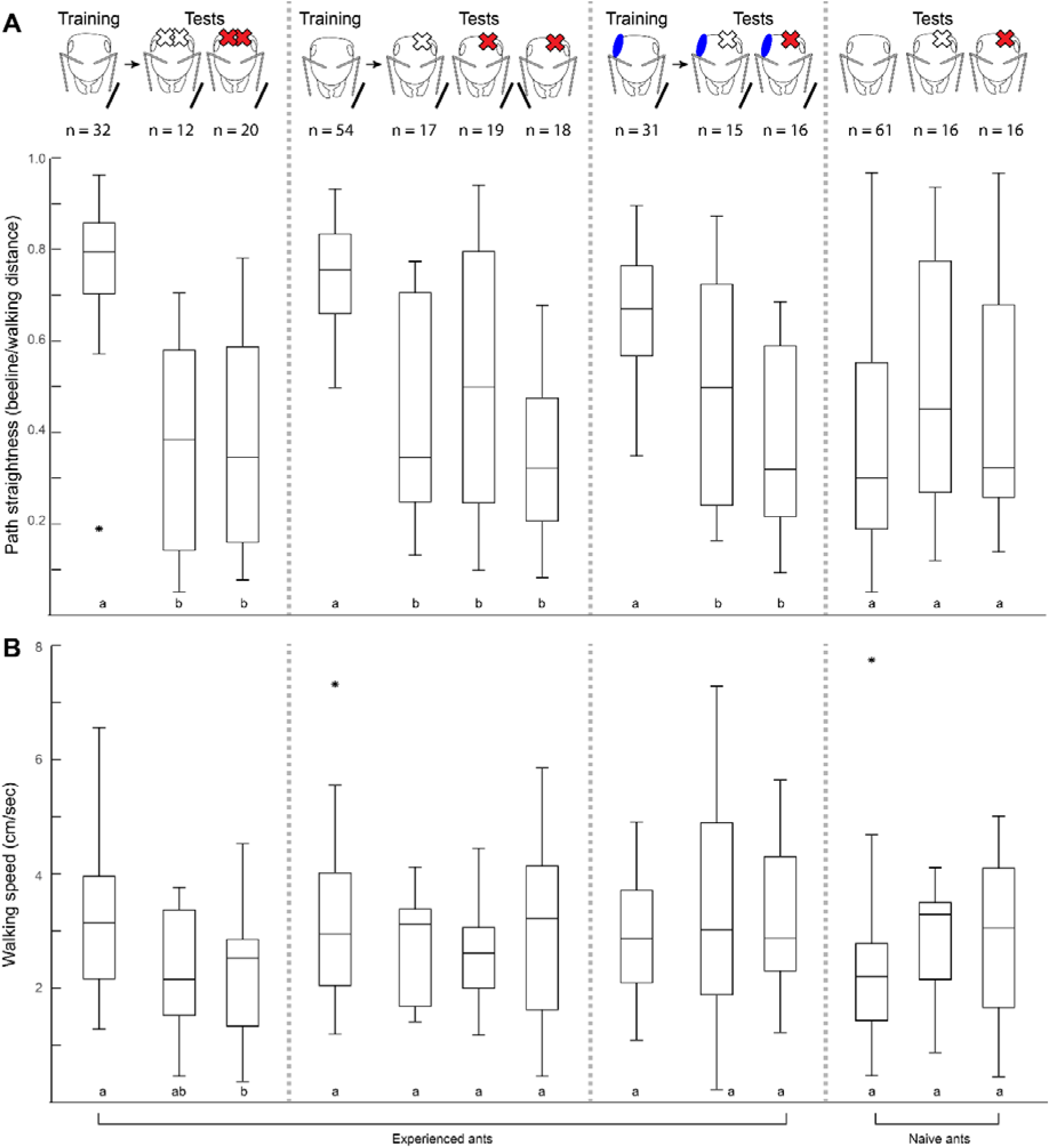
Path straightness and walking speed of ants with different physiological manipulations. Different physiological manipulations are shown in the schematic ant heads. Injection sites are shown as X (white, control injection; red, injection of local anaesthetic) and the black bar to the side of the ant’s head represents the side on which the visual cue would appear as the ant progressed to the feeder. Number of ants is shown at the top. Lowercase letters indicate significant differences (p<0.05) between the different groups of ants (Kruskal Wallis test with Dunn’s post hoc and Bonferroni corrections). Groups with the same letters are not significantly different. **(A)** Path straightness for ants from the different groups. **(B)** Walking speed from ants of the different groups.

**Figure S3:**
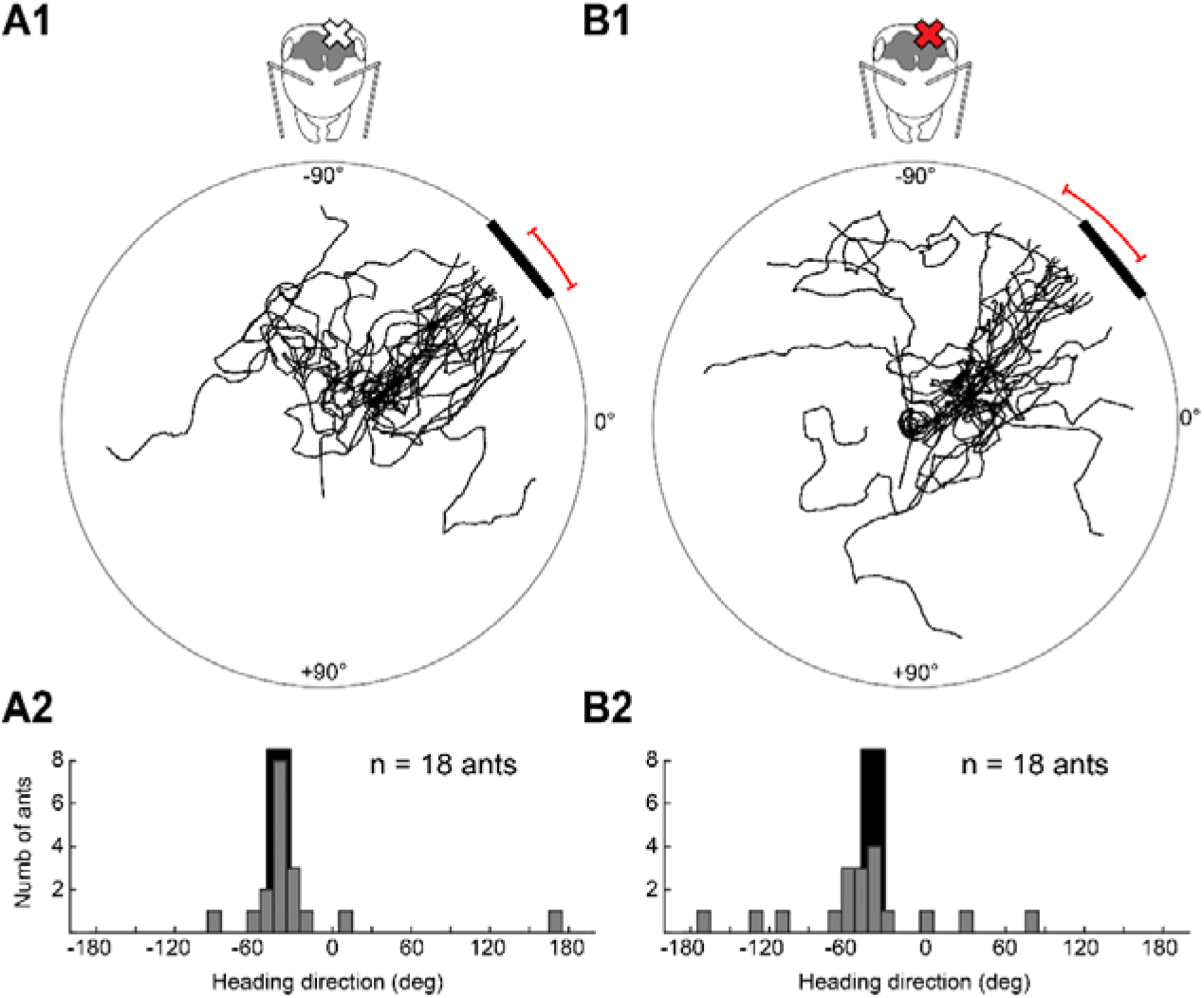
MB calyces are not required for innate attraction to visual cues. **(A)** The innate response of naïve ants was tested in the behavioural arena shown in Figure 1A (see also Figure S1A for paths of intact naïve ants). Naive ants had unilateral injections of the control solution into the MB calyx (X, injection site). For path plot and histogram description see Figure S1. **(B)** Same experimental procedure as in (A) but these ants had the local anaesthetic injected into the MB calyx.

## Acknowledgements

We would like to thank Scarlett Dell-Cronin, Sophie Atkinson and Jose Adrian Vega Vermehren for help with the experiments, Marta Rossi for help with the fluorescent dyes and Patrick Schultheiss for advice on chemical injections. The research was funded by the Biotechnology and Biological Sciences Research Council (Grant Number: BB/R005036/1).

## Author Contributions

Conceptualisation: CB, PG, JN, BWe

Methodology: CB, PG, JN

Formal Analysis: CB, BWe

Investigation: CB & BWo

Writing Original Draft: CB

Writing Review & Editing: CB, PG, JN, BWe, RG, BWo

Funding: PG, JN, BWe

## STAR METHODS

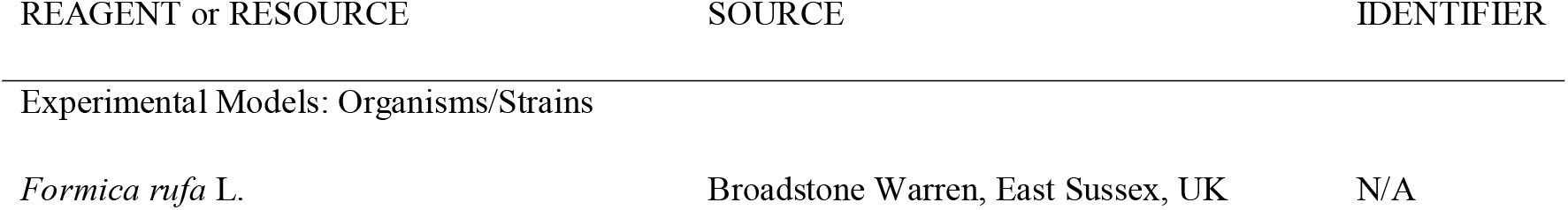
KEY RESOURCES TABLE

**KEY RESOURCES TABLE**
| REAGENT or RESOURCE | SOURCE | IDENTIFIER |
| --- | --- | --- |
| Experimental Models: Organisms/Strains |  |  |
| *Formica rufa* L. | Broadstone Warren, East Sussex, UK | N/A |

## Lead Contact And Materials Availability

Further information and requests for resources and reagents should be directed to and will be fulfilled by the Lead Contact, Cornelia Buehlmann.

## Experimental Model And Subject Details

Behavioural experiments were performed with laboratory kept wood ants *Formica rufa* L. collected from Broadstone Warren, East Sussex, UK. Ants were kept in the laboratory under a 12 h light: 12 h darkness cycle and constant temperature of 25-27°C. Ants were fed ad libitum with sucrose, dead crickets and water. During the experiments, food was limited to a minimum to increase the ants’ foraging motivation, but they had access to water all the time.

## Methodological Details

### Behavioural setup & experimental procedures

General experimental procedures for behavioural experiments followed those described previously [44]. Wood ant foragers learnt to find food at the edge of a circular platform (r = 60 cm) placed either 30° to the right or left of the outside edge of a single visual cue (Figure 1AB and Figure S1). After a few days of training, ants had fairly direct paths to the feeder and we started to perform brain lesions to investigate the role of the ants’ mushroom bodies during visual navigation. Chemical lesions were performed either bilaterally (Figure S1E1) or unilaterally (Figure S1E2) in the mushroom body calyces of well-trained individuals by either injecting a local anaesthetic coupled with a fluorescent dye (procaine hydrochloride and rhodamine in saline) or a control solution (rhodamine in saline). The ants’ navigational performance was recorded after the injection and the location of the lesion examined afterwards for each ant.

During training, individually marked ant foragers were taken from the nest and released in the centre of a circular platform (120 cm in diameter) that was surrounded by a cylinder (diameter 3 m, height 1.8 m) with white walls. Ants learnt to find a drop of sucrose on a microscope slide that, from the centre of the arena, was located 30° to the right or left, of the outside the edge of a 20° wide rectangle (height: 90 cm, width: 52 cm) placed on the inner wall of the surrounding cylinder. To remove possible olfactory traces, the surface of the platform was covered with white paper which was rotated after each round of training and was replaced between experiments. Ants performed approximately 10 group training runs before being trained individually. For individual training, ants were put separately into a 6.5 cm diameter, cylindrical holding chamber in the centre of the platform. The ant was released from the holding chamber by remotely lowering its wall. Once the ant had reached the sucrose slide and started to feed, the ant was transferred into a feeding box and the next ant was released. Ants were recorded using a tracking video camera (Trackit, SciTrackS GmbH) which provided the ant’s position on the platform every 20 ms. All individual training runs were recorded. When ants approached the feeder quite directly they were evaluated for testing. Ants were considered to be reliable and accurate if they approached the feeder directly on three consecutive training runs and only these ants were tested. In tests, ants were either chemically lesioned (see below for lesion and control injections) or an eye was occluded. In the eye occlusion test, one of the ants’ compound eye was covered with a layer of white enamel paint (Humbrol). The same eye capping procedure was applied for experiments where ants had one eye covered throughout training.

In addition to experiments with experienced ants, we also tested the role of the MBs in naïve ants. To do so, we selected active ants from the nest, performed the chemical lesion (either lesion or control injection) and recorded the ants’ innate response to the rectangular visual cue in the same arena as described above.

### Chemical brain lesions

Ants were immobilised on ice for 90 seconds and harnessed in a custom-made holder keeping their head fixed with plasticine while their body was able to freely move. Antennae were carefully held down with a pin. To access the ants’ MB calyces, a small window was cut with a piece of razor blade into the head capsule with four cuts: the lower cut was below the medial ocellus, the upper cut was above the two lateral ocelli, and the left and right cuts were between the left and right ocelli and the left and right compound eyes, respectively. Once the MB calyces were exposed, 0.5 nl of solution was injected uni- or bilaterally into the calyces. Injections were performed with a PV820 Pneumatic PicoPump (World Precision Instruments) connected to compressed air. Glass capillaries (Harvard Apparatus; 30-0035; 1.00 mm outer diameter, 0.78 mm inner diameter, 10 mm length) were pulled with a P-97 Micropipette Puller (Sutter Instrument) and then broken manually to a tip size of 10 μm. To get an injection volume of 0.5 nl with each capillary, we injected droplets of the solution into paraffin oil before each lesion to measure the droplet size produced by the capillary and adjusted the settings of the pico pump accordingly. Each capillary was only used once, i.e. two capillaries were prepared and calibrated for bilateral injections. After the solution was injected, the piece of head capsule was put back and fixed with a tiny droplet of super glue (Loctite, Power Flex). After surgery, ants were given 30 minutes for recovery before their behaviour was recorded.

Test ants were injected with either a control or test solution. For the lesioned ants, we used the transient and local anaesthetic procaine that selectively silences the neural activity in the MB calyces by reversibly blocking voltage-gated Na^+^ and other voltage-gated channels [11, 17–19]. Two stock solutions were prepared in advance and kept in the freezer for up to one month. Solution 1: A 40% procaine solution was prepared by diluting procaine hydrochloride (Sigma Aldrich; CAS 51-05-8) in ant saline (from [33]: 127 mM NaCl (CAS 7647-14-5), 7 mM KCl (CAS 7447-40-7), 1.5 mM CaCl_2_ (CAS 10043-52-4), 0.8 mM Na_2_HPO_4_ (CAS 7558-79-4), 0.4 mM KH_2_PO_4_ (CAS 7778-77-0), 4.8 mM TES (CAS 7365-44-8), 3.2 mM Trehalose (CAS 6138-23-4), pH adjusted to 7.0; all chemicals from Sigma Aldrich). Solution 2: 6 mM rhodamine B (fluorescent dye; Sigma Aldrich; CAS 81-88-9) was diluted in ant saline. During the experiments, solution 1 was diluted to 3 mM with saline to get the control solution. In order to get the lesion solution, solutions 1 and 2 were mixed in a 1:1 ratio to get a 20% procaine solution in 3 mM rhodamine. Lesion and control solutions were kept in the fridge for up to 3 days.

After the behavioural recordings, the ants’ brains were dissected in saline and imaged using a Nikon AZ 100 fluorescent microscope. Only ants with rhodamine-stained calyces were included in the analysis.

## Quantification And Statistical Analysis

The ants’ path directions were determined at r = 50 cm. Directionality of data was tested using the Ryleigh test for circular data [45]. To provide a plausible statistic for the heading direction while making minimal assumptions, for each condition we calculated a bootstrap distribution of the median using N=10,000 samples, and report the 95% confidence interval (2.5 and 97.5 percentile values) [46]. We would expect this interval to contain 0° if the ants in that condition were directed towards the feeder.

General walking speed and path straightness (index of straightness = beeline distance / path length) were calculated for paths from r = 11 cm to r = 50 cm (r being distance from the centre of the platform). Kruskal Wallis test with Dunn’s post hoc and Bonferroni corrections were used to compare ants from different groups.

## Data And Code Availability

Data will be available on the University of Sussex research repository (figshare).

## Additional Resources

